# Rapid evolution by spontaneous mutation increases genetic diversity facilitating plant population survival

**DOI:** 10.1101/497610

**Authors:** Henning Nottebrock, Mao-Lun Weng, Matthew T. Rutter, Charles B. Fenster

## Abstract

Using a mechanistic eco-evolutionary trait-based neighborhood-model, we quantify the impact of mutations on spatial interactions to better understand the potential effect of niche evolution through mutations on the population dynamics of *Arabidopsis thaliana*. We use 100 twenty-fifth generation mutation accumulation (MA) lines (genotypes) derived from one founder genotype to study mutational effects on neighbor responses in a field experiment. We created individual-based maps (15,000 individuals), including phenotypic variation, to quantify mutational effects within genotypes versus between genotypes on reproduction and survival. At small-scale, survival is enhanced but reproduction is decreased when a genotype is surrounded by different genotypes. At large-scale, seed set is facilitated by different genotypes while the same genotype has either no effect or negative effects. Mutations may provide a mechanism for plants to quickly evolve niches and may drive competition, facilitation and selection with profound consequences for future population and community dynamics.

## Introduction

A complete understanding of population and community dynamics requires linking intraspecific genetic diversity with spatial ecological interactions (Bolnick et al. 2011, Genung et al. 2011). Evolutionary mechanisms – mutation, drift, gene flow and selection – are responsible for intraspecific genetic variation contributing to both ecological structure and species diversity (Hart et al. 2016). These community properties impact ecological mechanisms such as competition and facilitation (Whitlock 2014). Competition and facilitation in turn affect population and community dynamics that depend on variation in demographic rates (Chase et al 2002, Solivers et al 2015). Demographic rates are influenced by spatial interactions of genetic variation and this variation may contribute to species coexistence: genotypes can hinder or favor the survival of each other by using similar or different resources with negative or positive consequences for coexisting species (Hart et al. 2016, Hausch et al. 2018). Intraspecific genetic variation contributing to individual trait variation and environmental adaptability may promote species coexistence by both increasing habitat heterogeneity and altering competitive hierarchies (Violle et al. 2012, Ehlers et al. 2016). However, intraspecific genetic variation may hinder species coexistence when intraspecific genetic variation is diminished by competition between individuals (Hart et al 2016). To fully understand the causes of coexistence mechanisms, we need to integrate intraspecific evolutionary mechanisms with intraspecific ecological mechanisms underlying spatial species interactions (Bolnick et al. 2011, Ehlers et al. 2016).

The number of studies exploring eco-evolutionary dynamics and their potential feedbacks on population and community dynamics has dramatically increased over the last two decades (Shefferson and Salguero-Gomez 2015). An important current issue in community and ecosystem genetics research is determining the relevance of intraspecific genetic variation and genetic differentiation (divergence) to ecological and evolutionary processes at the community and ecosystem level (Genung et al. 2011, Pujol et al. 2018). Quantifying intraspecific trait variation defining the fundamental niches of species is an important link between ecology and evolution (Violle and Jiang 2009). Spatial structure is another important component of realistic eco-evolutionary dynamics, such that in spatially structured populations, selection is determined by the interplay between demographic and genetic structures (Lion et al. 2011). Demographic structure describes the spatial distribution of individuals through birth, death and migration resulting in spatial patterns, while genetic structure describes the spatial distribution of genotypes. The amount and spatial pattern of genetic variation may constrain evolution of traits influencing competitive ability of individuals (Wilson 2014). Moreover, competitive abilities depend on species’ niches. The strength of competition is determined by how much individual niches overlap with each other (Hutchinson 1957, Holt 2009) and how long coexistence has occurred (Connell 1980).

Species niche breadths are reflected by individual trait variability and intraspecific and interspecific genetic and spatial interactions driven by environmental conditions. Thus, intraspecific trait variability shaped by intraspecific genetic variation can influence ecological mechanisms driving variation among individual persistence (Lankau 2009). Only when the spatial relationship between traits and demographic parameters is explicitly described can robust hypotheses about the effects of individual variation on competitive outcomes be accurately formulated (Hart et al. 2016). Especially in controlled common garden environments, genetic variation in one species can have predictable and heritable effects on associated communities and ecosystems (Carr and Dudash 1995; Whitham et al. 2003, 2006; Johnson & Stinchcombe 2007; Bailey et al. 2009; Johnson, Vellend & Stinchcombe 2009). Hence, coupling evolutionary genetics with community ecology may advance our understanding of species interactions and population and community dynamics (Baron et al. 2016), especially for populations and communities that suffer environmental change (Bellard et al. 2012).

An important gap in our understanding of the link between evolutionary and ecological processes is the nearly complete absence of data quantifying how rapidly genetic variation governing within species competitive hierarchies evolves (Hausch et al. 2018). An important source of novel population genetic variation is mutation. Therefore, we aim to investigate the link between evolutionary genetics and spatial ecological interactions of mutation accumulation lines (MA lines) in the model plant organism *Arabidopsis thaliana* (Brassicaceae) to advance our understanding of the genetic origins of population and community dynamics, contributing to a fuller understanding of the maintenance of biodiversity. We take advantage of 25th generation *A. thaliana* mutation accumulation (MA) lines that were planted under field conditions in years 2004 and 2005 with spatial records of each individual each year (e.g., Rutter et al. 2010; 2012, Rutter et al. 2018) (Fig. 1).

**Figure 1.**
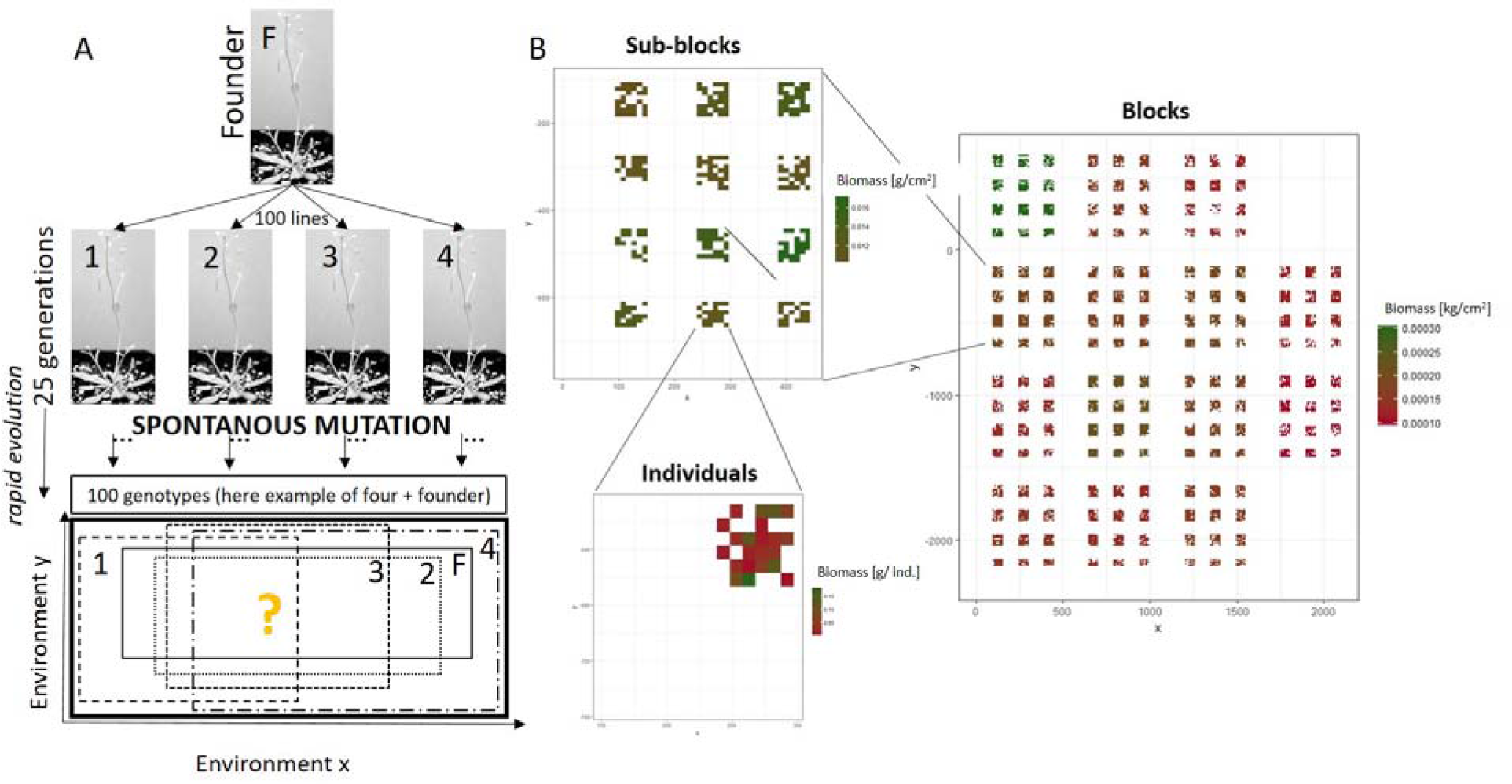
(A) 100 Mutation accumulation lines (here example of four MA-lines) of *Arabidopsis thaliana* derived from a single founder (Columbia) after 25 generations with a mutation rate of ~ 0.7 per generation (Weng et al. 2019). Each MA line has a unique genotype that differs from the founder. Fundamental niche of individual genotypes (different dashed and dotted lines) and founder genotype (black line) are distributed within the hypothetical fundamental niche of a population of *A. thaliana* (bold black line). Axes represent niche axis (resource-use, environmental factors, etc.). (B) Planting design of *A. thaliana* in a natural field experiment showing spatial biomass variation of individual survivors in 2004.

Here we focus on the neighborhood effects of MA lines on focal plants from the same MA line (within genotypes) or different MA lines (between genotypes) to quantify the evolution of intraspecific genetic variation through the accumulation of spontaneous mutations. We implement trait-based neighborhood models (Nottebrock et al. 2017a, Nottebrock et al. 2017b, Lachmuth et al. 2017) to understand ecological mechanisms of within and between MA lines genetic variation to examine eco-evolutionary dynamics (Lion 2018). Specifically, we ask: 1) Do spontaneous mutations rapidly introduce enough genetic variation among MA lines to influence competition and facilitation of *A. thaliana*? 2) Do spontaneous mutations of MA lines contribute to competition, facilitation and selection promoting differential MA line survival and reproduction at different spatial scales? 3) How may competitive and facilitative effects potentially determine population and community dynamics?

## Methods

MA lines are generated from a single nearly homozygous individual founder and cultivated via limited effective population number. In the case of *A. thaliana*, this occurs through single seed descent, resulting in Ne= 1. Thus typical MA line cultivation results in an unbiased sample of mutation effects ranging from deleterious to advantageous, although lethal mutations are excluded (Lynch and Walsh 1998). Each MA line accumulates independent spontaneous mutations. After the propagation of a set of MA lines through multiple generations, the genetic differences among the MA lines and between those lines and the founder reflect the input of mutation. Significant MA line effects for multiple traits, including performance and trait measures, were found under both field and greenhouse conditions (Rutter et al. 2010; Roles et al. 2016, Rutter et al. 2018). Each of the MA lines in our experiment is fixed for an average of 20 different sequence level mutations, single nucleotide mutations (SNMs) and indels combined (Ossowski et al 2010, Rutter et al. 2012, Weng et al. 2019).

### MA lines and field experiments

We used survival and seed set data of *A. thaliana* MA lines and the founder as assessed in field experiments in 2004 and 2005 from Rutter et al. (2010, 2012 and 2018) planted in a randomized design (Fig. 1). Rutter et al. (2010, 2012) planted seedlings of 100 MA lines and the founder at the four-leaf stage, approximately two weeks post germination, into a secondary successional field at Blandy Experimental Farm (BEF) in Virginia (39°N, 78°W). Each of the 100 MA lines was used to found up to five sublines to minimize biases due to maternal effects introduced by the specific location within the greenhouse. We founded six sublines from each of the six lines representing the premutation founder genotype. In 2003, subline plants were used to generate all seed utilized in all field experiments. In each planting, 7504 individuals were planted, 7000 individuals of 100 MA lines (70 replicates per MA line, 14 replicates per subline) and 504 individuals of the founder (14 replicates per subline). The planting environment corresponds to a spring ephemeral life-history, where plants germinate and complete the life-cycle in the spring. At the time of planting, vegetation was scant but present. By harvest, the *A. thaliana* individuals were dwarfed by naturally occurring vegetation.

The plot was arranged in 14 spatial blocks with each containing 12 subblocks (Fig. 1B) (total plot area approximately 35 × 25 m). Each block included one seedling from each subline and in total 7504 individuals. We used the spatial information of each individual within the described design and created a raster of all plant individuals with R packages (raster, maps, maptools). We used individual-based maps neighborhood matrices with exact spatial and trait information of each genotype. If all five sublines did not produce enough seedlings to distribute in all blocks, seedlings from other sublines of the same MA line were overrepresented in blocks to maintain the same overall number of plants per MA line. Plants dying within the first 3 days of transplant (about 50 plants) were considered to have died from transplant shock and were replaced with another plant from the same MA line. Plants were censused weekly for survival. Plants were harvested by late May, by which time they had senesced. In 2004 a total of 5915 individuals including 394 founders survived. In 2005, a total of 4506 individuals survived including 302 founders. Plants were oven dried and biomass was measured. All fruits produced by each plant were counted and in combination with seed production used as the measurements of seed set. For our analysis, we used measurements of two response variables representing an important part of the life history of *A. thaliana* to calculate direct neighborhood effects on: a) the survival rate of plant individuals and b) individual seed set from plants that survived and produced fruits.

### Statistical Analyses

We use eco-evolutionary trait-based neighborhood models that include intraspecific genetic variation and phenotypic variation expressed by plant trait biomass to analyze competition and facilitation between individuals of *A. thaliana*. We analyzed plant survival rate and focal seed set measurements from years 2004 and 2005 separately. For each year, we considered all individuals as focal plants in the analysis based on individual-based neighborhood matrices to analyze 1) rate of plant survival to reproduction and 2) seed set of those plants that survived to reproduction. We analyzed neighboring plants to focal plants in a radius of 80 cm (small-scale) or 200 cm (large-scale) of a given focal plant as two spatial scales in the neighborhood analyses (Fig. 1) to quantify selection, competition or facilitation between plants depending on their genotypic and phenotypic variation.

We used extensions of linear mixed models (package Ime4, Bates et al. 2014) in R ver. 3.3.3 (www.r-project.org) to conduct neighborhood analyses of focal seed set and survival. We assumed binomial errors for the analyses of plant survival and Poisson errors for analyses of seed set. The mixed models described interactions among plants by including neighborhood indices as explanatory variables at two spatial scales in separate models. Neighborhood indices are spatial density effects of surrounding neighborhood plants that affect focal seed set and survival. For each plant, we used the Euclidian distance between the focal plant and the neighboring plants to compute response effects of intra-(same genotype) or inter-genotypic (different genotype) neighbors in a given radius around focal plants (Nottebrock et al. 2017). Moreover, we used a neighborhood index that accounts for the decline of neighbor effects with distance from the focal plant (Uriarte et al. 2010) and summed the amount of biomass from all individuals in a radius of 80 cm or 200 cm respectively by a Gaussian interaction kernel (Lachmuth et al. 2018, Nottebrock et al. 2017, Damgaard 2004). We used random effects of block and subblock to correct for environmental variation between local heterogeneous conditions. Importantly, we correct for between MA line effects by including MA lines as a random effect. In addition, including a random slope of biomass on each random intercept corrects for the intraspecifc phenotypic variation depending on local conditions of plant focal individuals. Moreover, the weighted neighborhood density by plant biomass accounts for environmental variation between neighboring plants. Our model parameters and a detailed model description of subblock and block models can be found in SI supplementary text (Supplementary Material).

Neighborhood matrices of all individuals (individual-based maps) were used to analyze the effect of intra- and inter-genotypic neighbors on survival and focal seed set with spatial interaction kernels of neighborhood (plant biomass) density. By incorporating different genotypes and phenotypic variation, we can quantify how important genetic variation is for neighborhood models and if the phenotypic variation explains spatial interactions between individuals. We assume the consequences of genetic differences to be larger between MA lines than between any MA line with the founder. This is a valid assumption since each MA line differs from the other by approximately 20 + 20 = 40 mutations, while any two replicates within a MA line will differ by one generation, ≤ 2 mutations. Thus, we simulated line effects from parameters derived from MA lines as random effects with the R package ‘merTools’ and the function ‘plotREsim’ in R 2018.

We used the trait values of neighbors to calculate trait-based neighborhood indices including plant biomass as a trait (Goldberg & Fleetwood, 1987; Goldberg & Landa, 1991; Cahill et al., 2005). We fitted eco-evolutionary trait-based neighborhood models at two different spatial scales for response variables (survival and seed set) for each of the two and both years. To address our objectives, we first analyzed models with differential effect in which intra-genotypic neighbors (within all MA lines and founder) had a different effect on survival (A1, A2, Table 1) and seed set (B1, B2, Table 1) than inter-genotypic neighbors at small-scale (s) and large-scale (I). In addition, we analyzed models with neutral effects on survival (A1, A2, Table 1) and seed set (B1, B2, Table 1) that included total neighbor density without the split between intra-genotypic and inter-genotypic neighbors at small-scale (s) and large-scale (I). To this end, all models were fitted with two separate neighborhood indices that were calculated from intra- and inter-genotypic neighbors. To justify the inclusion of individual plant biomass as trait-values for interacting plants in the model, we used AlCc to compare the models with and without the trait-proxy (Burnham and Anderson 2002). We found that all models perform better including biomass as a trait-proxy (ΔAICc > 2). All eco-evolutionary trait-based neighborhood models contained random effects of subblock nested in block at block scale and subblock scale on the intercept, MA line identity on the intercept and the focal trait-value (plant biomass) on the slope. Additionally, because direct environmental variables were not measured during the field experiments, we included in each model the individual’s biomass to correct for environmental conditions for spatial autocorrelation. All variables are scaled and centered to assure comparability between predictor variables. Models of differential and neutral effects for 2004 and 2005 (Table 1, A1-A2, B1-B2) are fitted at small-scale (80cm scale) and at large-scale (200cm scale). Hereafter, the 80 cm scale models are referred to as “small-scale” models and the 200 cm scale models are referred to as “large scale” models. Neighborhood indices, intra- and intergenotypic variation and total variation of biomass density are included as inverse density variables (1/1+density). We compared models of differential and neutral effects through likelihood ratio tests (LRTs).

**Table 1.**
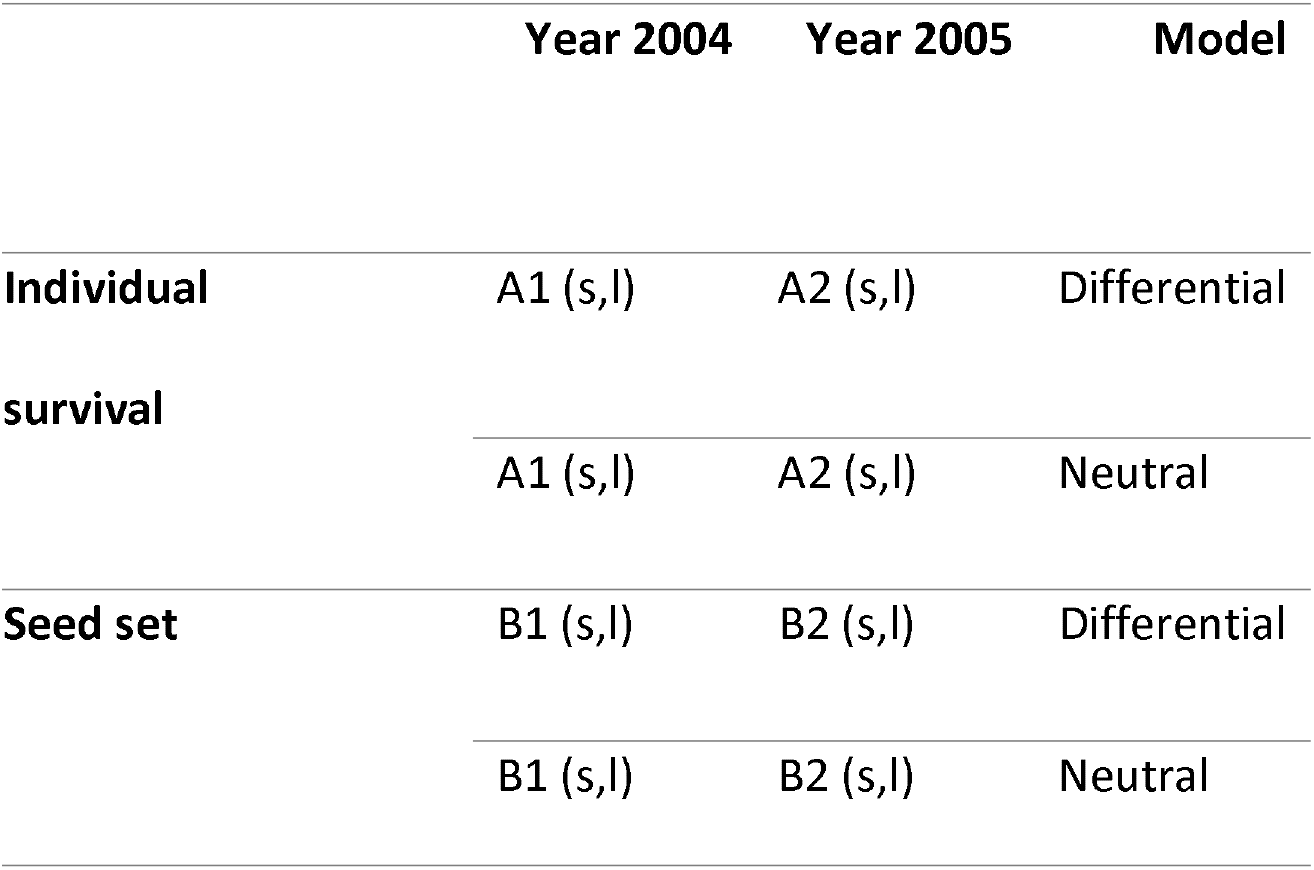
Table of all different eco-evolutionary trait-based neighborhood models including intraspecific variation to test our three objectives of density effects on survival and seed set of *A. thaliana*. Differential models split the neighbor responses into within genotypes and between genotypes. Neutral models describe the overall neighbor response without differentiating between genotypes. All models include random effects of plant biomass on the slope, subblock or block and MA-line on the intercept (see Fig. 1 for design), and respectively depending on small (s) or large-scale (I) (for more details see method section).

## Results

Focal plants surrounded by plants of the same genotypes varied from 0 to 6 individuals at small-scale (sub-block) and 5 to 6 individuals at large-scale (block). After determining that eco-evolutionary trait-based models perform better when including plant biomass as a trait proxy to calculate trait-based neighborhood indices we found that all models including plant biomass perform better than models including only neighborhood density (ΔAIC > 2, Supplementary Material S2).

### Weighted neighbor effects of biomass density on genotypes for plant survival

At small-scale, survival rate is larger when surrounded by inter-genotypic than intra-genotypic neighbors for both years 2004 and 2005 (model A1 and A2; Table 2 and Fig. 2a). This difference of survival rate between intra- and inter-genotypic plants indicates that inter-genotypic neighbors select for genotypic diversity and show stronger competitive effects on the survival of intra-genotypic neighbors at small-scale. This finding is demonstrated by the superior performance of the differential model relative to the neutral model (year 2004: LRT: *χ*^2^_1df_ = 5.21, p < 0.05; year 2005: LRT: *χ*^2^_1df_ = 14.73, p < 0.001). The differential and neutral models of survival at large-scale have only non-significant effects (Table 2). In addition, comparing the AlC we found that models at large-scale perform worse than at small-scale for both 2004 and 2005 (Table 2). We therefore will only discuss small-scale effects on survival. The distance kernel (alpha) at small-scale shows that neighboring plants of different genotypes reduce plant survival by 50% at 53 cm in 2004 and 48 cm in 2005.

**Figure 2.**
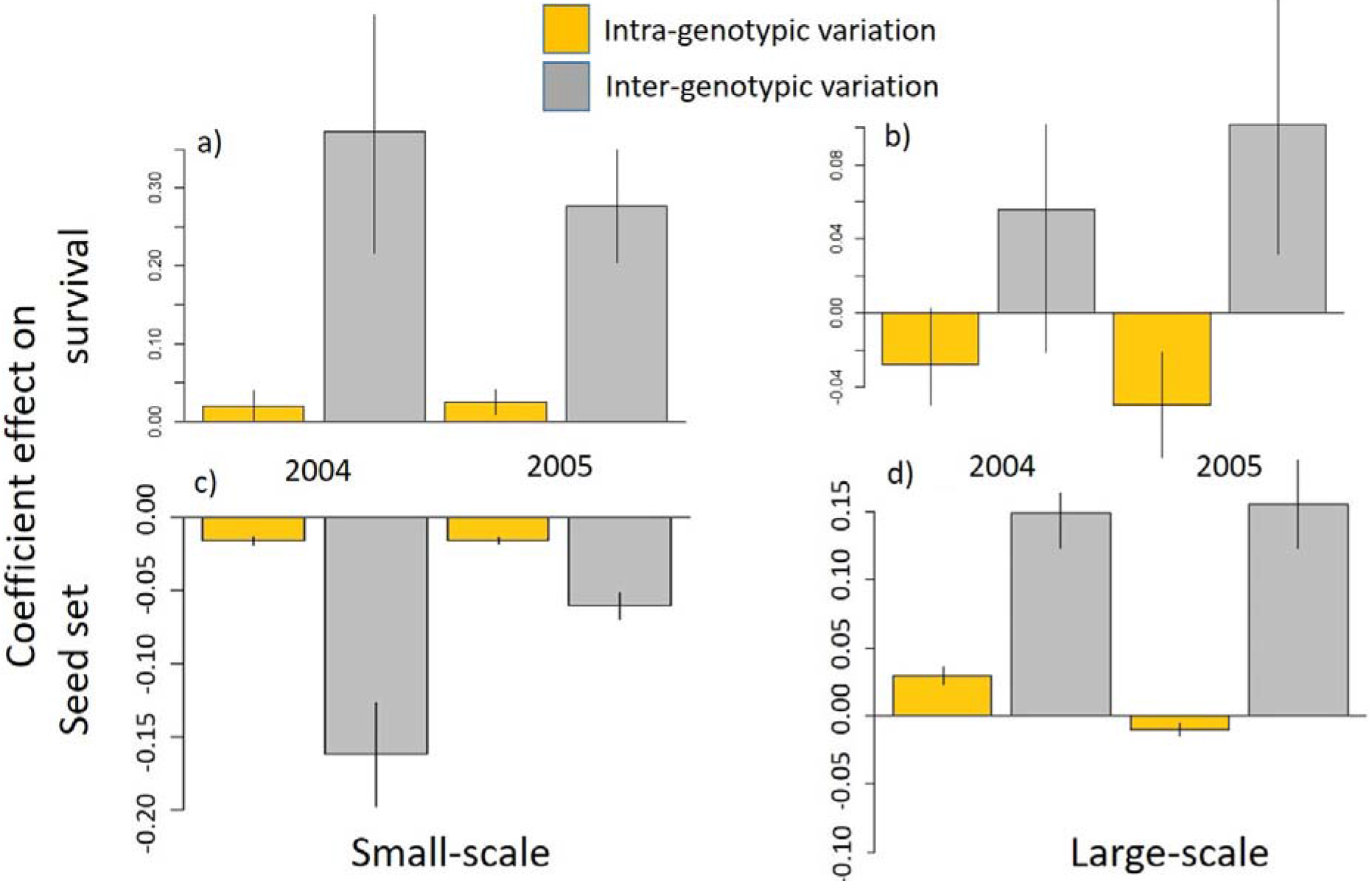
The barplots show standardized coefficient effect sizes and associated standard errors of eco-evolutionary trait-based neighborhood models at small-scale (a, c) and large-scale (b, d) for all MA-lines for the survival model (a, b) and the seed set model (c, d). Intra-genotypic effects (yellow) and inter-genotypic effects (grey) are presented for 2004 and 2005. Coefficient effect sizes indicate reduction or an increase of seed set depending on biomass density of intra- or inter-genotypic neighbors. In addition, coefficient effect sizes indicate a reduction of survival of intra or inter-genotypic neighbors.

**Table 2.**
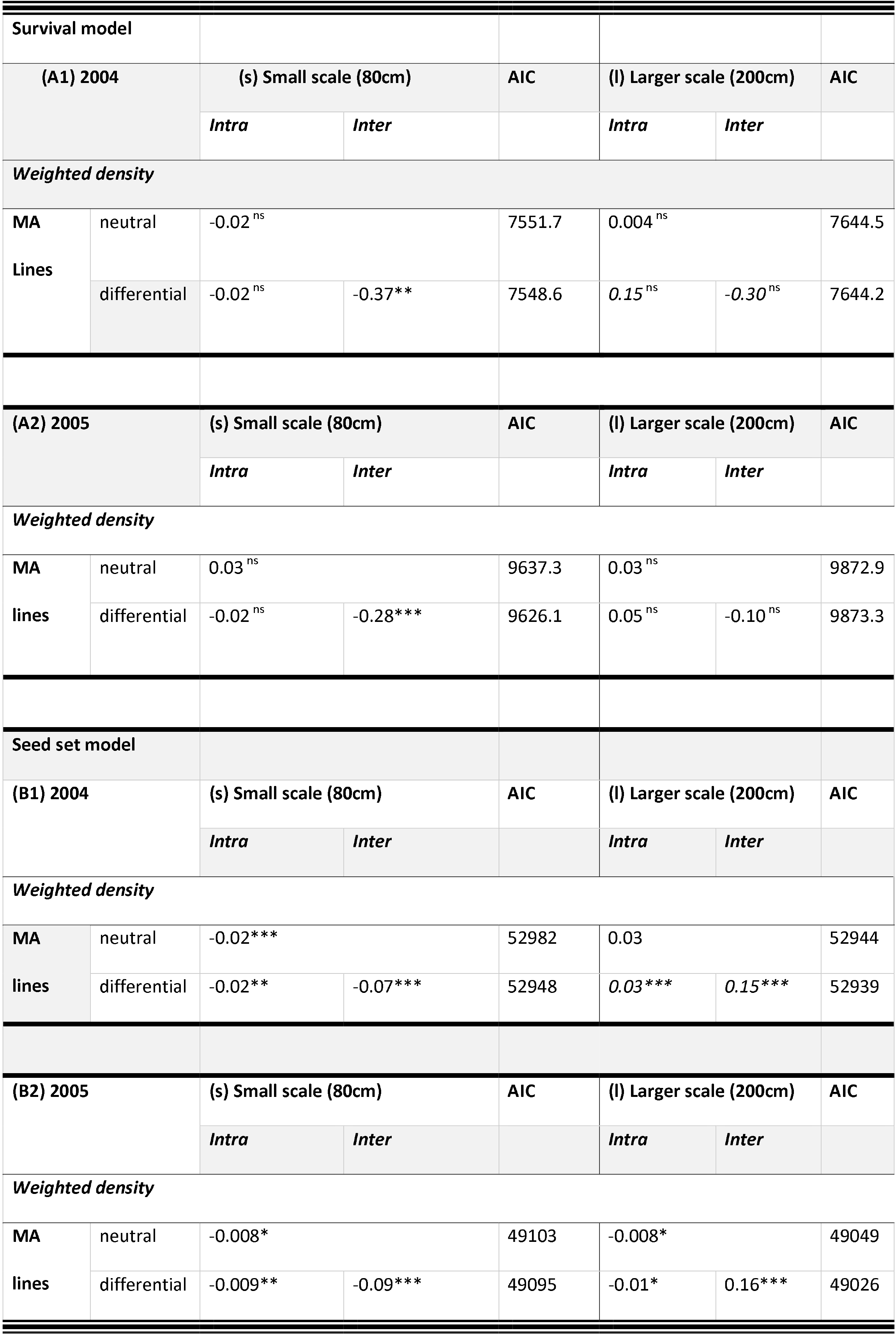
The coefficient and AlC from eco-evolutionary trait-based neighborhood models of the *Arabidopsis thaliana* field experiment in the years 2004 and 2005. Models include intraspecific genetic variation defined as intra-genotyptic (within genotypes) and inter-genotypic (between genotypes) variation. Models are shown at two different spatial scales: weighted subblock density (80cm radius) and weighted block density (200cm radius). All scales include variables of neutral effects (absence of intra- and inter-genotypic variation (total sum of all neighboring genotypes)) and differential effects (presence of intra- and inter-genotypic variation (split of neighboring genotypes in intra- and inter-genotypic genotypes). Thus, neighbor identity is split into intra- and inter-genotypes as differential effects or are combined as neutral effects, which includes all neighbors. P-value levels <0.05*, <0.005**, <0.001*** and >0.05^ns^.

### Weighted neighbor effects of biomass density on genotypes for seed set

At both small and large scales, the differential model performs better than the neutral model: at small scales (year 2004: LRT: *χ*^2^_1df_ = 52.48, p < 0.001; year 2005: LRT: *χ*^2^_1df_ = 17.37, p < 0.001; respectively) and at large scale (year 2004; LRT: *χ*^2^_1df_ = 679.44, p < 0.001; year 2005: LRT: *χ*^2^_1df_ = 21.17, p < 0.001; respectively). The small-scale model of seed set that included weighted biomass densities demonstrates the reduction of seed set is larger when surrounded by intergenotypic than intra-genotypic neighbors for both years 2004 and 2005 (model B1 and B2; Table 2 and Fig. 2a). The difference between intra- and inter-genotypic plants indicates that inter-genotypic neighbors have a stronger competitive effect on seed set than intra-genotypic neighbors at small-scale. In contrast, at large-scale seed set increases when surrounded by inter-genotypic neighbors but decreases when surrounded by intra-genotypic neighbors for both years 2004 and 2005 (model B1 and B2; Table 2 and Fig. 2b). Comparing the AlC between small and large-scales, models at large-scale for 2004 and 2005 perform better (Table 2).

### Genotype effects on plant survival and seed set

The distance kernel at small-scale in 2004 indicates a reduction of seed set (competition) whereas at large-scale are consistent with an increase of seed set (facilitation). Neighboring plants from all genotypes of *A. thaliana* at small-scale reduce 50% of focal seed set at 23 cm and at large-scale facilitate 50% of focal seed set at 86 cm. The estimation of genotype effects (MA lines and founder) simulated as conditional means and expressed as odds ratios show hierarchical orders of MA line and founder competitive effects at small-scale (Fig. 3). We found no significance of simulated competitive effects of all MA lines and founder calculated from survival and seed set models at small-scale (ANOVA, p > 0.1, Fig. 4a). In addition, we found no significant correlation between competitive effects of MA lines and founder at large-scale. In addition, we did not find selective effects of MA lines and founder on the survival at small-scale (ANOVA, p > 0.1, Fig. 4b). However, we found strong correlation between competitive effects of MA lines and founder at small-scale and large-scale indicating a genetic trade-off (ANOVA, F=460.07, P< 0.0001, Fig. 4c).

**Figure 3.**
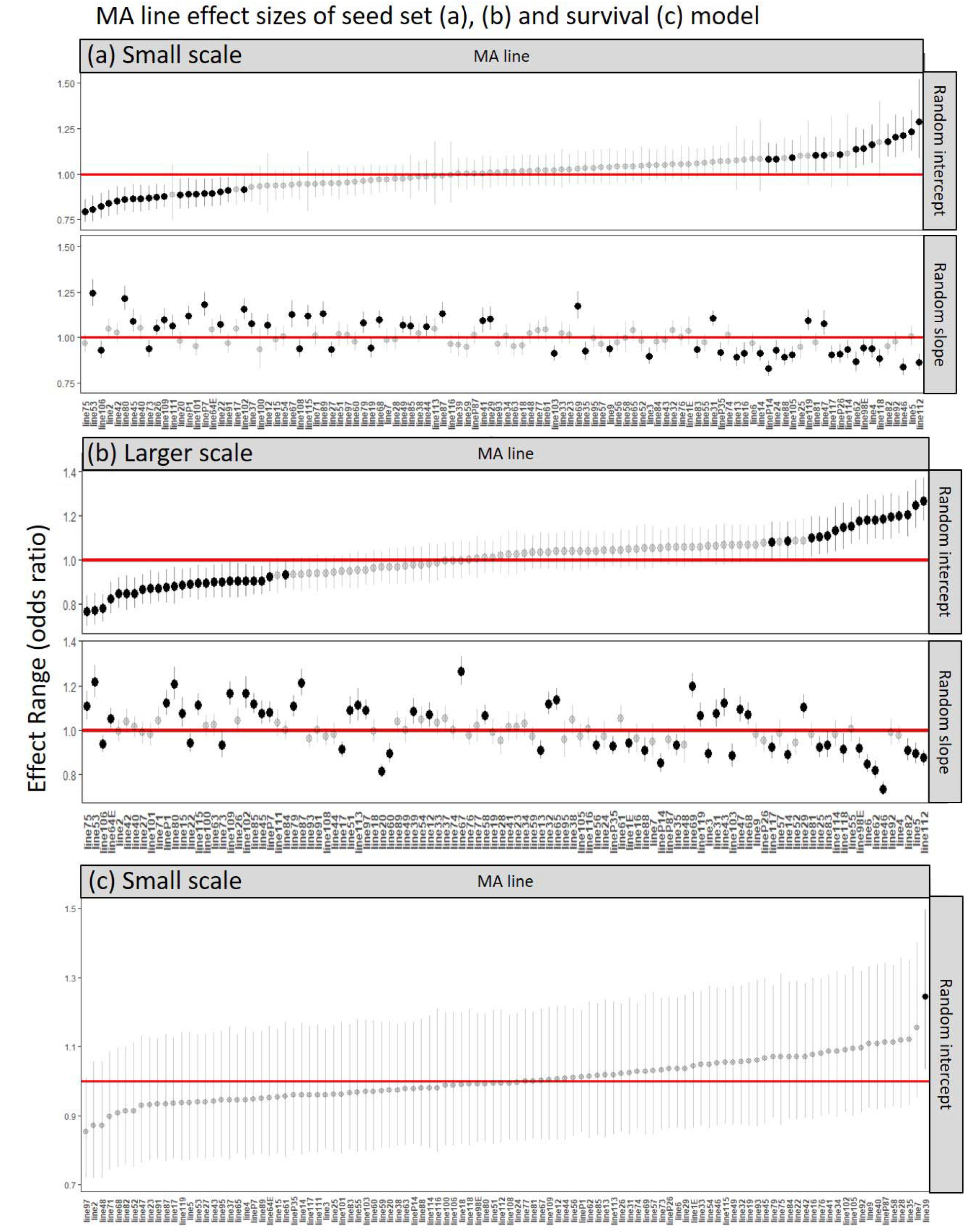
The effect range on seed set (a), (b) and survival (c) computed from eco-evolutionary trait-based neighborhood models shown as odds ratios for year 2004 (for 2005 see Supplementary Material), (a) Shows the hierarchical competitive order of MA lines and founder for the seed set model including range effects of each MA line and founder as random intercepts and the trait proxy biomass as random slopes at small-scale and (b) at large-scale, (c) Shows the hierarchical selective order of MA lines and founder for the survival model including range effects of each MA line and founder as random intercepts at small-scale

**Figure 4.**
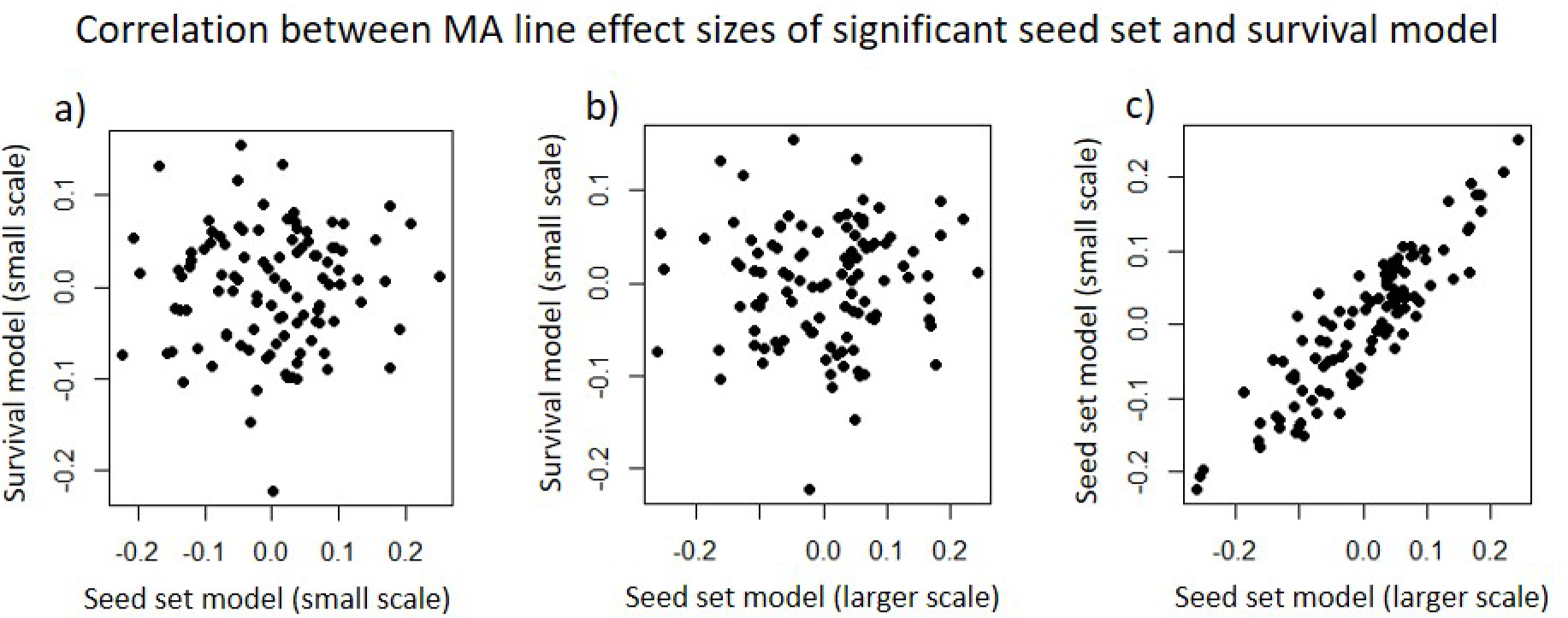
Correlation of the simulated conditional mean represented as odds ratio of same genotypes (MA lines) reducing or increasing the effect sizes of the seed set model and of the survival model at small-scale or large-scale for 2004, respectively. For year 2005 see supplementary material. Each point represents different genotypes of *A. thaliana*. Negative values means lower effect sizes on the survival or seed set and positive values means increasing the effect sizes on the survival or seed set. a) Relationship between survival and seed set model, b) Relationship between survival and seed set model, c) Relationship between small-scale survival and large-scale seed set model. Since we are plotting effect sizes, the positive correlation in panel c reflects a trade-off between competition and facilitation (see results).

## Discussion

We are able to quantify individual spatial interactions between and within genotypes (MA lines and founder) using a mechanistic eco-evolutionary trait-based neighborhood model. We demonstrate intraspecific genetic variation due to spontaneous mutations can shape competitive abilities of genetically different individuals of *A. thaliana*. Notably, these differences arose in just 25 generations of mutation and in the absence of natural selection. Many of the mutations differentiating the MA lines also appear in nature, contributing to *A. thaliana* genetic polymorphism (Weng et al., 2019). Plant survival is higher when surrounded by inter-genotypic neighbors supporting genetic diversity. Effects on focal seed sets were reversed between small and large scales; inter-genotypic neighbors have stronger negative (competitive) or positive (facilitative) effects than intra-genotypic neighbors on focal seed sets at small-scale or large-scale, respectively. Moreover, at small-scale, competitive effects of different MA lines have similar impacts on survival and seed set. At large-scale, the intra-genotypic effect on survival turns into less competitive effects on seed set. In contrast, different genotypes show stronger facilitative effects within the population at large-scale, maybe because plants are more competitive against other species. Below we discuss how these scale effects provide insight on community assembly and potential coexistence.

### Selection, Competition and Facilitation

Because of stronger statistical signals of effect sizes between genotypes than within genotypes, niche variation may result in niche expansion or reduction (Ehlers et al. 2016). However, these competitive effects only occur between plants that survive to reproduction and affect seed-set. The neighbor effect on plant survival suggests kin recognition despite our randomized field experiment that likely reduced the spatial autocorrelation of genotypes (Hamilton 1964, Kubisch et al. 2013). Seed-dispersal is limited in space because the *A. thaliana* fruit do not dehiscently explode as they do in related species (Hofhuis and Hay 2017), although a seed bank ensures some dispersal through time (Falahati-Anbaren et al. 2014). The neighbor effect increasing genetic diversity of *A. thaliana* populations can have a positive effect on the coexistence of competing species (Vellend 2006). If we assume that the increase of genetic diversity due to spontaneous mutations of *A. thaliana* affects the extent of individual plant niches, at large-scale the inclusion of other plant species should result in stronger competitive effects of *A. thaliana* with other plant species (Hausch et al. 2018). We demonstrate that *A. thaliana* individuals increase their competitive abilities with an increase of genetic diversity, perhaps due to the expansion of their individual niches due to more variation in plant traits.

Antagonistic interactions between genotypes on different spatial scales may result in genetic trade-offs when mutations are advantageous or deleterious, which over many generations could provide a strong stabilizing force maintaining both species and genetic diversity in this system and promote coexistence (Lankau 2008). Thus, spontaneous mutation could also provide an additional evolutionarily stabilizing effect on community dynamics. Our study demonstrates the potential for a fundamental evolutionary process, mutation, to have profound consequences for community structure. When *A. thaliana* is rare, selection would favor genotypes that compete well and enhance the population’s survival relative to that of its interspecific competitor (Lankau 2009). In contrast, when *A. thaliana* is common and the interspecific competitor is rare, selection would favor *A. thaliana* genotypes that are good intraspecific competitors. This trade-off may result in a decrease in the interspecific competitive ability of *A. thaliana*, effectively increasing the competitor’s fitness relative to that of *A. thaliana* (Lankau 2009, Lankau and Strauss 2007). However, the genetic trade-off between intra-and interspecific competitive abilities due to mutation remains unknown because our dataset only includes performance of *A. thaliana* individuals without quantifying the surrounding intraspecific environment (Chesson 2000, Adler et al. 2007).

### Competitive Hierarchies of MA lines and Founder lines

Our result show scale dependent competition and facilitation (Nottebrock et al. 2017b) providing further evidence that in a relative short evolutionary time scale spontaneous mutations may change the competitive hierarchies between founders and specific MA lines due to advantageous or deleterious mutations (Rutter et al. 2010). In contrast to Masclaux et al. 2010, we found differential responses to similar genotypes vs different genotypes in *A. thaliana* depending on the neighbor effect on the plant survival or seed set. However, the interaction between similar genotypes and different genotypes seems to depend on the strength of the competitive abilities of the accessions (this study). Comparisons between MA lines and founder demonstrate that competitive hierarchies follow different orders and competitive effects occur at different life-history stages at different spatial scales. Eco-evolutionary processes might reflect spatial selection for diversified genotypes because of niche evolution and individuals of different genotypes that survived to reproduction have stronger competitive abilities (Ehlers et al. 2016). Moreover, the hierarchical order of competitive effects between MA lines and founders shows strong variation.

### Eco-evolutionary dynamics

Populations can adapt evolutionarily to their environment on a time scale equivalent to that of ecological processes and affect present day species interactions, i.e., coexistence (Fussmann et al. 2007). Yet, sustained and rapid climate change could deplete genetic variance faster than it can be replenished by mutation (Fournier-Level et al. 2016). In our study, we show that only 25 generations are enough to influence present day plant interactions of *A. thaliana* in the presence of other plant species. Possible effects of mutations on trait mechanisms might reflect the increased seed set of *A. thaliana* at large-scale, because of the presence of another plant species allowing the establishment and growth of *A. thaliana* (Ehlers et al. 2016). Many traits can influence competition and facilitation between individuals of different genotypes (e.g. Chapin et al. 1993; Caradus and Woodfield 1998; Hausmann et al. 2005). Intraspecific competition between *A. thaliana* individuals tends to be higher than between interspecific competitors (Lankau and Strauss 2007, Van Dam and Baldwin 1998; Tiffin 2002, Strauss and Irwin 2004). Such spatial mechanisms might be described by the scale difference of competition at small-scale and facilitation at large-scale between different *A. thaliana* genotypes. Competition and facilitation are therefore dependent on the extension or reduction of individual niches, which is based on the ability to advance or distract traits due to spontaneous mutations. However, we are not able to directly link our results to benefactor or antagonistic plants, because we focused in our experiment on *A. thaliana* densities and fitness components. Nevertheless, genotypic-specific interactions are well studied. A number of studies of *A. thaliana* MA lines demonstrate mutation effects on plant traits such as leaf weight, flowering time, trichome density, number of leaves at bolting, duration of vegetative and flowering period (Camara and Pigliucci 1999, Rutter et al. 2010, Stearns and Fenster 2016). It is conceivable that the increase in variance of these traits through spontaneous mutations could contribute to the evolution of competitive hierarchies.

### Consequences for population and community dynamics

Competition between genetically different individuals of *A. thaliana* at small-scale and facilitation at large-scale has important implications for population structure (Cahill et al. 2005). Plants may cooperate by competing less, or act selfishly by competing more (Dudley et al. 2013). However, kin recognition might be the result of kin selection when occurring in a heterogeneous environment with various genotypes (Hamilton 1964). Especially, at large-scale, we show that different genotypes facilitate seed set that stabilize the population dynamics by increasing the performance of neighboring plant individuals (Latzel et al. 2013, Castellanos et al. 2014). Thus, selection for diversified genotypes might reveal a coexistence mechanism, because mutations at a short time scale (25 generations) are able to alter the competitive hierarchies of *A. thaliana* individuals of different genotypes. These genotypes might expand or reduce their niches and can therefore act more competitive against strangers (Ehlers et al 2016). Moreover, variation in competitive ability among genotypes due to mutation can lead to intransitive competitive hierarchies at a small-scale, and allow coexistence of competitors at large-scale when there is no single dominant competitor (Vellend and Geber 2005, Taylor and Aarssen 1990, Laure et al. 2017). This might also reflect the genetic trade-off between competitive abilities of MA lines of small-scale competition and of large-scale facilitation (Timan 2004). Low competition between *A. thaliana* individuals at small-scale transmits into low facilitation between different species at larger scale. In turn, strong competition within *A. thaliana* at small-scale results in weaker competition between different species at large-scale. Thus, the constant input of mutation to genetic variation of competitive hierarchies suggests that genetic variation is likely never a limiting factor in the evolution of those traits that influence plant community dynamics.

### Implementation of eco-evolutionary trait-based model for population and community ecology

In ecology, two primary contrasting models are crucial to population and community dynamics for understanding the maintenance of biodiversity: the niche and neutral theories of ecology (Fisher and Mehta 2015). The niche theory claims that niches are determined by the difference of abiotic or biotic ecological processes e.g. competition or facilitation (Hutchinson 1957, Bruno et al. 2003, Colwell and Rangel 2009) whereas the neutral theory predicts that ecological processes are neutral and are solely modified by stochasticity (Hubbel 2005). Although it is still under debate which model predicts the dynamic properties of communities and determines the maintenance of biodiversity (Violle et al. 2017), both models miss the importance of evolutionary mechanisms and individual variation influencing population and community dynamics (Violle et al. 2012). The difference of competition and facilitation between or within genotypes demonstrates that evolutionary mechanisms that shape individual variation could change hierarchies of intra-specific interactions in natural plant populations over a short evolutionary time scale. Our results are concordant with the niche theory stressing the changing of competitive hierarchies among phenotypes in a given environment (Levine and HilleRisLambers 2009, Nottebrock et al. 2017). Moreover, selection and trait evolution favor population survival of *A. thaliana* in the wild, which stresses the importance of combining spatial ecological and evolutionary mechanisms for our understanding of population and community dynamics.

## Conclusion

Rapid evolution of *A. thaliana* due to spontaneous mutation alone has profound consequences for population and community dynamics. Competition of *A. thaliana* individuals is genotype-dependent and *A. thaliana* is not a single weak competitor among co-occurring plant species (Ehlers et al. 2016, Soliveres et al 2017). Moreover, the discovery of the underlying eco-evolutionary nature of competition in *A. thaliana* supports a shift from species-based to individual-based community ecology. This would lead to a more predictive ecological theory (Violle et al 2017). We show that including the intraspecific genetic variation and phenotypic variation as separate aspects to a trait-based neighborhood model increases the predictive power to understand population and community dynamics. In particular, non-neutral intraspecific processes may determine species coexistence, because genetic diversity is promoted by having stronger competitive abilities at small-scale and stabilizing population survival at large-scale (Clark et al. 2010). Additionally, our study demonstrates higher genetic diversity increases population survival due to rapid evolution with implications to forecast the fate of species and functional diversity in response to environmental changes (Violle et al 2012). Depending on the time of environmental changes, species may adapt to environmental change by shifting their fundamental niches (Clark 2010). The combination of intraspecific genotypic variation and spatial interactions might advance our understanding of community dynamics, especially of rapid evolution (Koch et al. 2014, Turcotte and Levine 2016). Incorporating genetic variation and the eco-evolutionary process for determining standing levels of genetic variation will provide a better understanding of species interactions underlying the maintenance of biodiversity.

